# Quantitative differences in developmental profiles of spontaneous activity in cortical and hippocampal cultures

**DOI:** 10.1101/009845

**Authors:** Paul Charlesworth, Ellese Cotterill, Andrew Morton, Seth G. N. Grant, Stephen J. Eglen

## Abstract

**Background:** Neural circuits can spontaneously generate complex spatiotemporal firing patterns during development. This spontaneous activity is thought to help guide development of the nervous system. In this study, we had two aims. First, to characterise the changes in spontaneous activity in cultures of developing networks of either hippocampal or cortical neurons dissociated from mouse. Second, to assess whether there are any functional differences in the patterns of activity in hippocampal and cortical networks.

**Results:** We used multielectrode arrays to record the development of spontaneous activity in cultured networks of either hippocampal or cortical neurons every two or three days for the first month after plating. Within a few days of culturing, networks exhibited spontaneous activity. This activity strengthened and then stabilised typically around 21 days *in vitro*. We quantified the activity patterns in hippocampal and cortical networks using eleven features. Three out of eleven features showed striking differences in activity between hippocampal and cortical networks. 1: Interburst intervals are less variable in spike trains from hippocampal cultures. 2: Hippocampal networks have higher correlations. 3: Hippocampal networks generate more robust theta bursting patterns. Machine learning techniques confirmed that these differences in patterning are sufficient to reliably classify recordings at any given age as either hippocampal or cortical networks.

**Conclusions:** Although cultured networks of hippocampal and cortical networks both generate spontaneous activity that changes over time, at any given time we can reliably detect differences in the activity patterns. We anticipate that this quantitative framework could have applications in many areas, including neurotoxicity testing and for characterising phenotype of different mutant mice. All code and data relating to this report are freely available for others to use.

## Introduction

During development, many parts of the nervous system generate patterns of spontaneous activity. These patterns of activity are thought to be instructive in the assembly of neural connectivity, for example by driving activity-dependent mechanisms [1]. To date, most recordings of spontaneous activity have been *in vitro*, although recent *in vivo* studies using calcium imaging also report the presence of patterned spontaneous activity [2, 3]. *In vitro* recordings are typically made with multielectrode arrays (MEAs) which contain at least 60 electrodes. These recordings allow us to assess activity at a range of levels from the single-unit to the network. Beyond its relevance for understanding how activity might guide development of the nervous system, spontaneous activity recordings have also been used as an assay for network performance in applied settings, like neurotoxicity screening [4].

In recent years there has been significant interest in measuring the developmental patterns of spontaneous activity in networks cultured from neurons in control and experimental conditions [4, 5, 6, 7, 8]. Although many properties of spontaneous activity have been reported, we do not yet have a systematic sense of how these features change across development, or which features of neural activity are useful at describing the observed patterns of activity.

To address both these questions, we have cultured two types of network on MEAs and recorded their activity every 2–3 days up to around one month post-plating of neurons onto the array. In the first type of network, we cultured hippocampal neurons taken from embryonic mice. The second type of network was created using exactly the same protocol except with neurons dissected from cortex. Recordings of spontaneous activity from both types of network were quantified using eleven different features at the level of individual electrodes, pairs of electrodes, or the entire array. We found that hippocampal networks tend to generate more regular bursting activity, including theta bursts, and more correlated activity than the corresponding cortical networks at the same age.

## Results

### Development of spontaneous activity

Within seven days of culturing neurons on MEAs, spontaneous activity can be reliably recorded (Fig. 1) from both hippocampal and cortical networks. As development progresses, we find that the firing rate increases, and that the frequency of bursting increases. To quantify these differences, we have used a range of measures (Fig. 2) to assess the activity at a single-electrode level, pairwise, and at the level of the entire network. All of these measures are defined in the methods.

**Figure 1.**
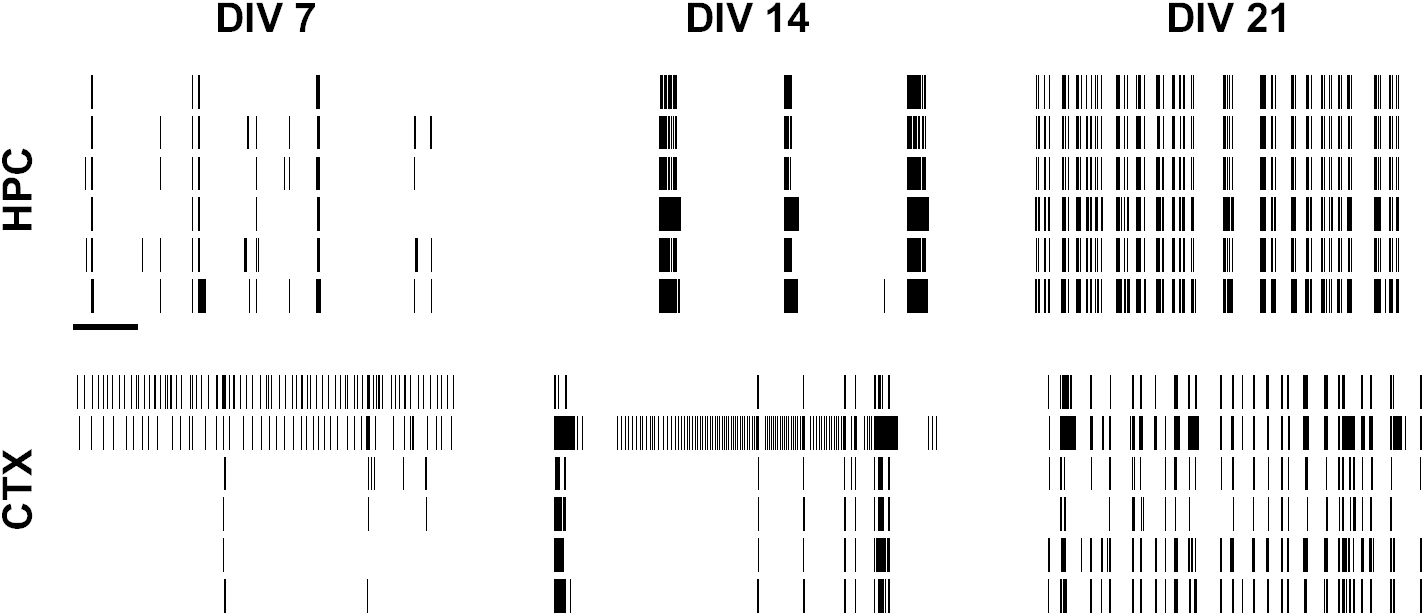
Examples of spontaneous activity in developing hippocampal (HPC; top row) and cortical (CTX; bottom row) cultures. Each column represents one day in vitro (DIV). Within each raster plot, one row represents the spike train from one electrode; six (out of typically 59) electrodes are shown. Scale bar for all rasters is 10 s.

**Figure 2.**
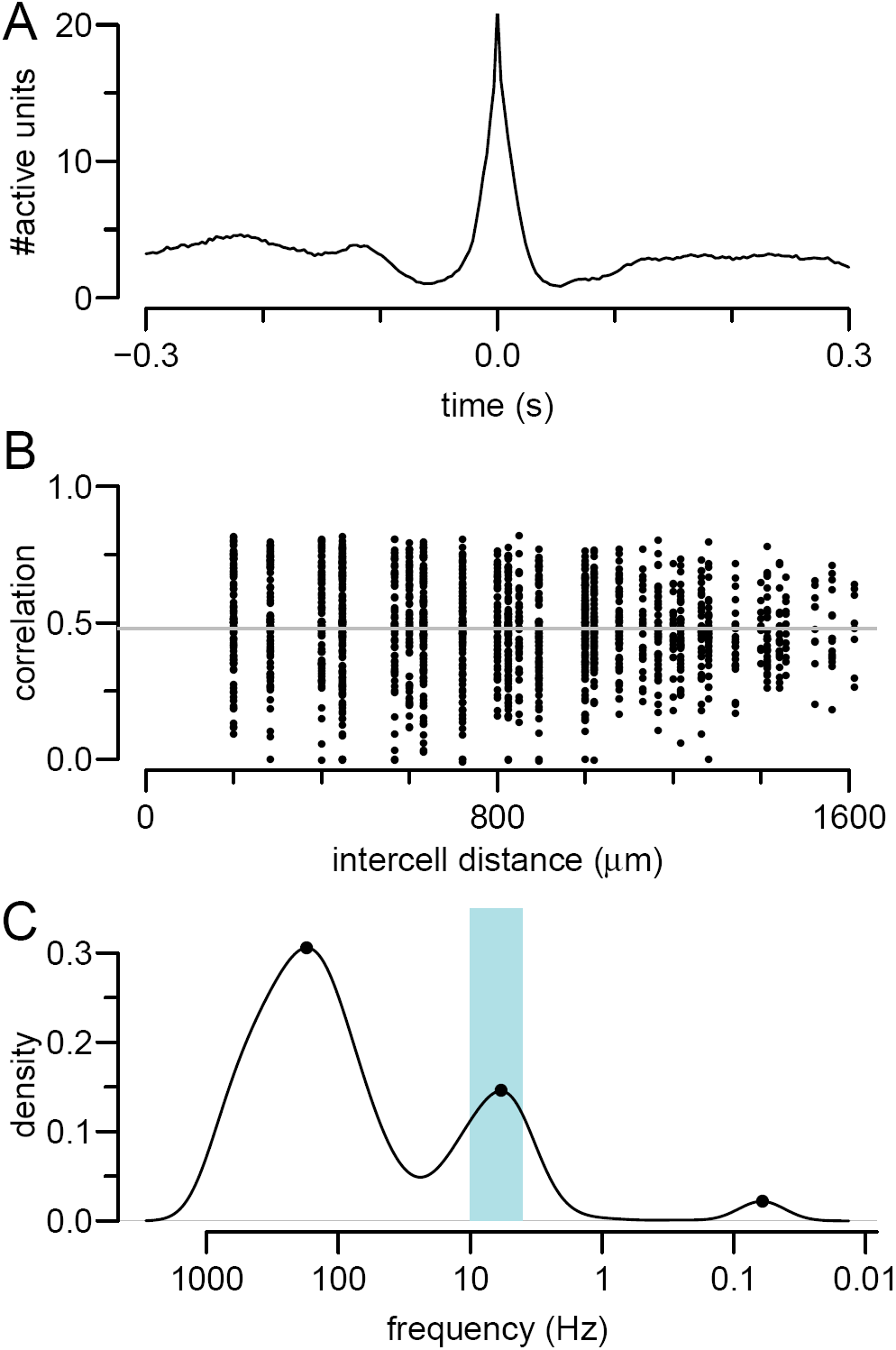
Examples of features calculated for each recording. The hippocampal recording from 14 DIV in Figure 1 was used as an example for this figure. A: mean network spike. B: pairwise correlation calculated using the spike time tiling coefficient. As there is weak dependence on distance, we take the mean (grey solid line). C: detection of theta bursting on an electrode with a firing rate close to the median activity on the array.

*Overall firing rates* During development, there are slight, statistically-significant differences in firing rates, with median firing rates being slightly higher for hippocampal networks, but overall there are no key differences at maturity (Fig. 3A).

**Figure 3.**
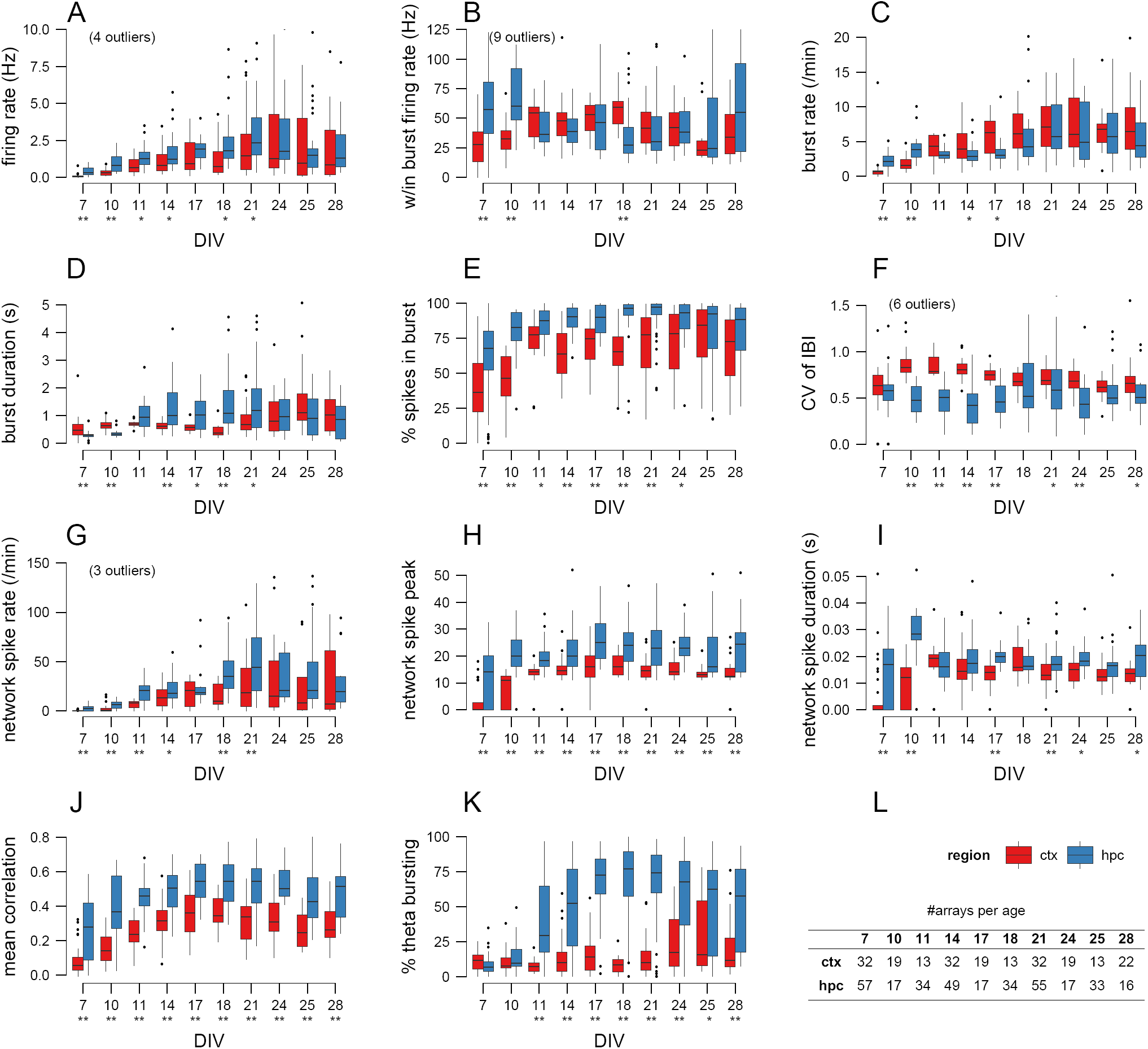
Characterisation of spontaneous activity in hippocampal and cortical networks. Panels A–K shows the values of one feature (named on the y axis) as a function of age. Boxplots show the median and interquartile range, with whiskers extending out the most extreme values within 1.5 times the interquartile range. Individual points outside this range are regarded as outliers and drawn as points; in a few cases these outliers are not drawn to keep the y-axis within a meaningful range. Underneath each age, stars denote significant difference of median values for cortical and hippocampal networks at either 0.05 (*) or 0.01 (**) level (Mann-Whitney test, with p values corrected for multiple comparisons with false discovery rate method). L: number of arrays analysed at each age.

#### Bursting properties

Neurons typically fire in bursts, and are thought to be a reliable unit of neuronal information for functions such as coincidence detection and synaptic modification [9]. We find that bursts emerge around 7 DIV (days *in vitro*) and strengthen until about 14 DIV after which the bursting properties tend to stabilise. Among the bursting properties that we have measured, two factors seem to differentiate hippocampal and cortical networks. First, there is a higher fraction of spikes occurring within bursts for hippocampal networks (Fig. 3E), although the difference is no longer significant by 28 DIV. Second, the spike trains from hippocampal networks seem to be more regular, as indicated by the lower coefficient of variation for interburst intervals (CV of IBI) (Fig. 3F). The other burst-based measures that we calculated, namely within-burst firing rate (Fig. 3B), burst rate (Fig. 3C) and duration (Fig. 3D) show weaker differences between the two types of network.

#### Network activity

The previous measures analysed spiking data independently on each electrode. As a first approximation to assessing network activity, we used the concept of “network spikes” [10] to define the degree to which activity is coordinated across the entire array. At any time *t* we count the number of active electrodes; when this count exceeds a threshold, we say that a network spike has occurred. We measured three properties of these network spikes: their rate (per minute), their duration and their peak amplitude. Hippocampal networks tend to have more network spikes than cortical networks (Fig. 3G) and the network spikes involve more electrodes across development (Fig. 3H). The hippocampal network spikes tend to last slightly longer, although this is not consistent across development (Fig. 3I). Overall, this suggests that network activity tends to be stronger and more coordinated in hippocampal than cortical networks.

#### Pairwise correlations

As a further method to detect coincident activity on electrodes, we computed correlation coefficients for all possible pairs of electrodes on the array. For any pair of spike trains, we computed the spike time tiling coefficient, as this measure is particularly well-suited for relatively sparse spike trains [11]. For *N* (typically 59) electrodes on the array we compute *N*(*N* − 1)/2 correlation coefficients (i.e. ignoring autocorrelations) and plot them as a function of the distance separating the two electrodes (Fig. 2B). This technique has been used in studies of spontaneous activity in developing retina, and often reveals that correlations are distance-dependent, typically following a decaying-exponential profile [12]. However, we found that there is little, if any, distance-dependence upon the correlation coefficients (Fig. 2B), similar to that reported before [13]. We therefore decided to compute the average of all pairwise correlations.

Across all developmental ages, we find that the mean correlation is higher in hippocampal than in cortical networks (Fig. 3J). From 7 DIV to 14 DIV we see that the mean correlation becomes reliably stronger; after 14 DIV the correlations tend to stabilise.

#### Presence of theta bursting

The theta rhythm is a prominent 4–10 Hz oscillation measured in the hippocampus, and is thought to be involved in a range of neural functions [14]. We decided to examine whether our networks exhibited such oscillations by checking for peaks in the log interspike interval histogram in the range 0.1–0.25 s. Figure 2B shows an example of one electrode (recording from a 14 DIV hippocampal network) that exhibited theta bursting. Our approach was to measure the fraction of electrodes exhibiting theta bursting. Perhaps the most striking feature that discriminates hippocampal from cortical networks is the presence of theta bursting in hippocampal networks. Although only about 10% of electrodes in hippocampal networks are classified as theta bursting at 7 DIV, it is after 11 DIV that theta bursting is found on 50–75% of electrodes (Fig. 3K). By contrast, most electrodes in cortical networks do not detect theta bursting, except at 25 DIV.

### Discrimination of hippocampal and cortical networks

Each of the eleven features documented in Figure 3 shows that there are significant differences between hippocampal and cortical networks. However, given that the distributions of values can overlap and yet still be statistically significant (e.g. firing rate at 21 DIV; Fig. 3A), we cannot use individual features to reliably discriminate between the two types of network. We therefore used machine learning techniques to address two related questions:

1. Out of the eleven features, which are the most important for discriminating hippocampal versus cortical networks?
2. Given a recording of a network at a given age, is it possible to predict the identity (hippocampal or cortical) of the network?

We therefore translated each fifteen minute recording into an eleven-dimensional vector, with element *i* of the vector storing the value of feature *i* measured from the recording. This vector is described below as a *feature vector* of the recording.

#### Principal Components Analysis

If there is a consistent difference in the properties of hippocampal and cortical recordings, we would hope that the corresponding feature vectors cluster into two distinct regions. However, as these feature vectors are eleven-dimensional, we must first reduce their dimensionality to visualise them. Many such dimensionality-reduction techniques are available; we chose to use the best-known method, principal components analysis. Figure 4 shows the projection of the feature vectors at three different ages down into two-dimensional space. At 7 DIV there is significant overlap between the hippocampal and cortical recordings, which might suggest that is hard to discriminate between the two types of recordings; however at 14 and 21 DIV the recordings from the same cell type cluster and there is significant separation of the hippocampal and cortical recordings. The graphs underneath each scatter plot show the cumulative percentage of variance accounted for by the components. In each case, the first two principal components account for at least 60% of the variance.

**Figure 4.**
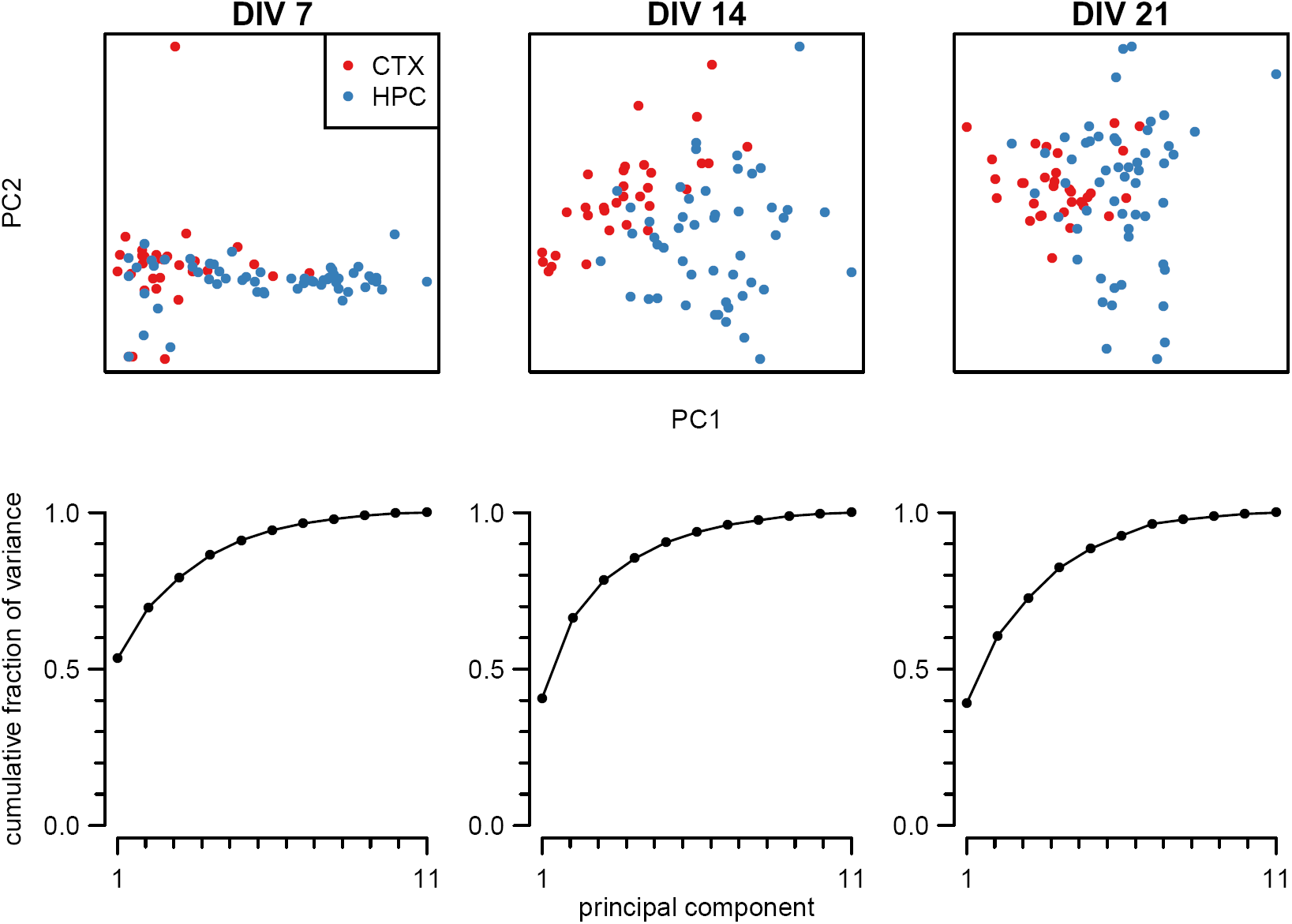
Principal components analysis of hippocampal and cortical feature vectors. Each column represents principal components analysis of the 11-dimensional feature vectors of all recordings at a given age (days in vitro). In the scatter plot, each point represents one recording projected down into the two dimensions that account for maximal variance and is coloured according to its cell type. Each graph shows the cumulative fraction of variance accounted for by the principal components.

#### Classification of recordings

The principal components analysis suggests that, especially at the latter ages, the feature vectors contain sufficient signal to discriminate between hippocampal and cortical networks. However, given the overlap between clusters, we next used classification techniques to quantify the degree to which the two classes of recording can be separated. We used two classification methods, detailed below. In both methods, 2/3 of the feature vectors at a given age are used to train a classifier to discriminate between the two types of recording. The remaining 1/3 of the feature vectors are then used as a test set to evaluate how well the classifier performs on data unseen during training.

We first used classification trees [15]. We built ten classifiers, one per age studied, to test whether the recordings could be grouped into cortical or hippocampal networks. We found that for any given age, the prediction accuracy of the trees was high — usually over 75% correct, depending on the age of the recording. (Performance would be 50% if there were no information to distinguish the two types of recording.) Once these trees had been built, we were able to interrogate them to find out which features were dominant in driving the classification of the network. At different ages, unsurprisingly, different features were dominant, but an overall trend clearly emerged when we averaged across developmental ages. Table 1 lists the features in decreasing order of their importance, along with their relative score (column 2).

**Table 1.**
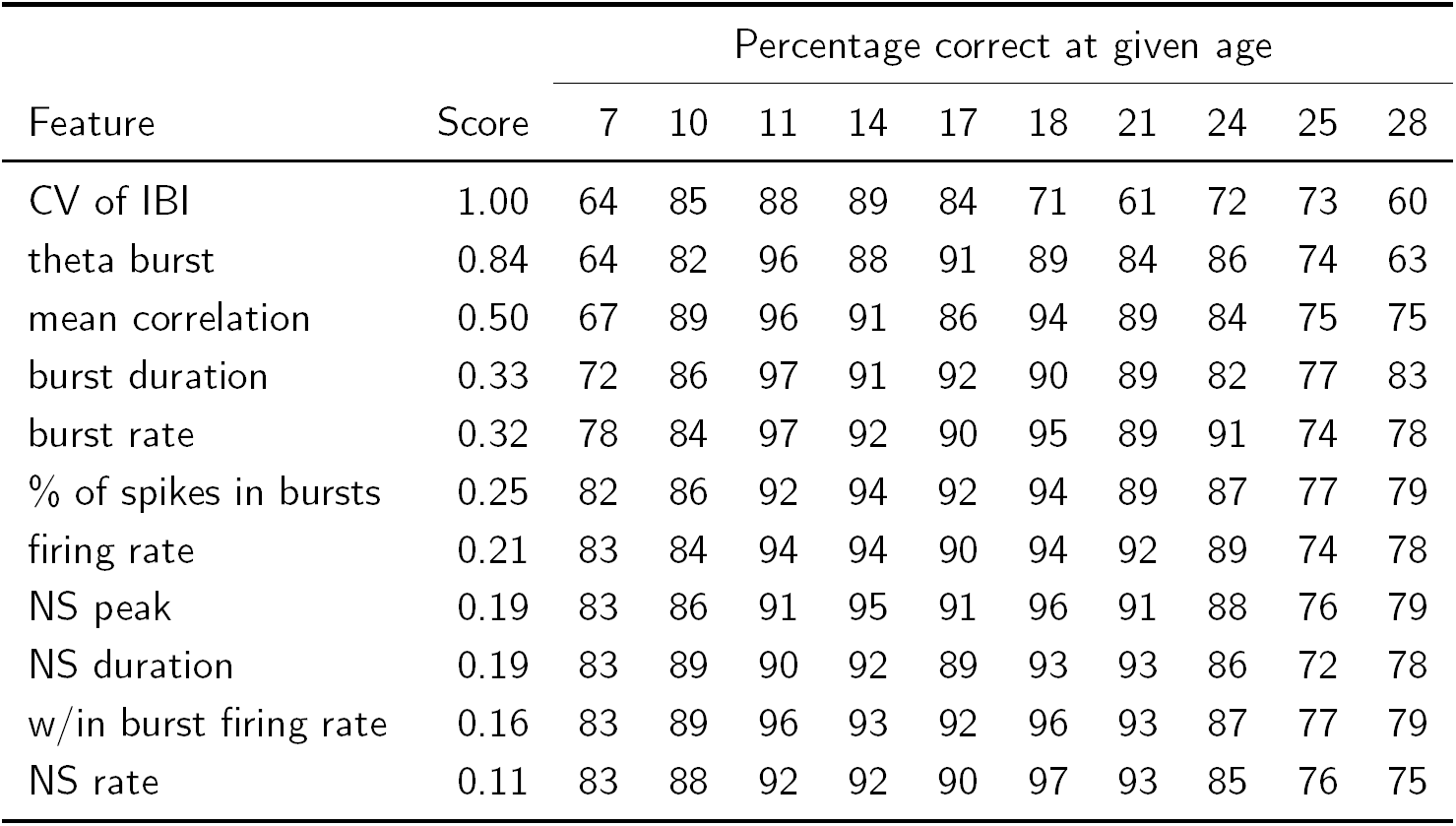
Classifier performance at discriminating cortical from hippocampal cultures. Features are listed in decreasing order of importance (score; column 2) normalised to the top score. The following numbers in each row *i* = 1 … 11 are the mean percentage of correct classifications at each age using the top *i* features.

| Feature | Score | Percentage correct at given age |  |  |  |  |  |  |  |  |  |
| --- | --- | --- | --- | --- | --- | --- | --- | --- | --- | --- | --- |
|  |  | 7 | 10 | 11 | 14 | 17 | 18 | 21 | 24 | 25 | 28 |
| CV of IBI | 1.00 | 64 | 85 | 88 | 89 | 84 | 71 | 61 | 72 | 73 | 60 |
| theta burst | 0.84 | 64 | 82 | 96 | 88 | 91 | 89 | 84 | 86 | 74 | 63 |
| mean correlation | 0.50 | 67 | 89 | 96 | 91 | 86 | 94 | 89 | 84 | 75 | 75 |
| burst duration | 0.33 | 72 | 86 | 97 | 91 | 92 | 90 | 89 | 82 | 77 | 83 |
| burst rate | 0.32 | 78 | 84 | 97 | 92 | 90 | 95 | 89 | 91 | 74 | 78 |
| % of spikes in bursts | 0.25 | 82 | 86 | 92 | 94 | 92 | 94 | 89 | 87 | 77 | 79 |
| firing rate | 0.21 | 83 | 84 | 94 | 94 | 90 | 94 | 92 | 89 | 74 | 78 |
| NS peak | 0.19 | 83 | 86 | 91 | 95 | 91 | 96 | 91 | 88 | 76 | 79 |
| NS duration | 0.19 | 83 | 89 | 90 | 92 | 89 | 93 | 93 | 86 | 72 | 78 |
| w/in burst firing rate | 0.16 | 83 | 89 | 96 | 93 | 92 | 96 | 93 | 87 | 77 | 79 |
| NS rate | 0.11 | 83 | 88 | 92 | 92 | 90 | 97 | 93 | 85 | 76 | 75 |

Out of the eleven features, three stood out. The most important was CV of IBI; this is a measure of firing regularity, which tends to be higher in hippocampal networks. Second, theta bursting is a key indicator (once it emerges at DIV 11) of hippocampal networks. Third, mean correlation is one of the top three features roughly half the time, and again is higher in hippocampal networks.

We chose classification trees as our first classification method primarily because of their simplicity (there are no free parameters) and ability to easily assess which features are driving the classification. Although its classification performance was good, we compared its performance against another common machine learning technique, namely Support Vector Machines (SVMs). We found that the SVM classifiers tended to result in slightly higher classification accuracy than the classification trees; e.g. when all 11 features were used, performance was 75–97% across ages (bottom row of Table 1).

Finally, given that our classifier trees provide us with a natural ordering of the importance of features, we asked how performance varied as we reduced the number of features that each recording is represented by. We found that performance remained high even as the number of features was gradually reduced (moving up through the rows of Table 1). It is clear however that multiple features are required for good classification; when only the single-most important feature is used (top row of Table 1), performance was only just above chance at some ages. However, with only three or four features, we obtained good performance across all ages.

In conclusion, the results from the classifiers tell us that three features of network activity (CV of IBI, theta bursting, and mean correlation) are strong predictors of whether a recording is from a hippocampal or a cortical network.

## Discussion

We have found that cultured networks of either hippocampal or cortical neurons generate spontaneous activity. These patterns of activity change during development and already by 7 DIV significant differences in their activity patterns begin to emerge. We have developed a quantitative framework for examining these activity patterns. By calculating eleven features of activity patterns, we can represent each recording of spontaneous activity as a point in (eleven-dimensional) feature space. When we examine recordings from any one given developmental age, we find that recordings from the same neuronal type (cortical or hippocampal) cluster in this feature space such that we can reliably discriminate between hippocampal and cortical networks.

Furthermore, out of the eleven features, we find that three are critically important for this classification in feature space. First, the CV of IBI was most important on average in driving the classification. Hippocampal spike trains tend to fire in bursts that are more regularly spaced than spike trains from cortical neurons. Second, after about 11 DIV most electrodes in hippocampal recordings detect theta-bursting, compared to a minority in cortical recordings. Third, the mean correlation between pairs of electrodes tends to be higher in hippocampal networks. These three measures are all relatively simple, and measure activity on either a single-electrode level or from pairs of electrodes. By contrast, although significant differences were found in the network spike measures, most importantly, the peak of network spikes, (Fig. 3F), these measures were not deemed to be critical in classification.

We have deliberately chosen simple features to characterise spiking activity to see if they suffice to discriminate between cortical and hippocampal networks. It is entirely likely that other more complicated measures of activity, particularly at the network-level, may also reveal clear differences between these two types of network [7]. For example, network connectivity measures have been used to explore differences in spontaneous activity patterns between mature hippocampal and cortical networks in a range of frequency bands [16]. However, the simple measures we have chosen here suffice to reliably differentiate the two types of network. Likewise, even higher classification performance may be possible with more complex machine learning techniques. However, our primary interest was to see whether in principle the feature space can be reliably separated with standard approaches [15]. Similar machine learning methods are not yet routinely used in analysing spontaneous activity, although see [17] for a recent example showing how singlecell activity could be classified as either *in vivo* or *in vitro*. Finally, with the advent of a new generation of higher density MEAs containing up 4096 electrodes [18], it is likely that there are much richer patterns of activity than we describe here.

We believe that our framework lends itself nicely to many applications, for example in neurotoxicity testing where spontaneous activity from a network is recorded whilst it is exposed to a particular compound [4]. By building up a representative feature space of recordings from compounds known to be either toxic or safe, our approach can be used to predict the toxicity of novel compounds. This idea builds upon earlier work where mean profiles of activity in each condition were used as simple classifiers [19]. More recently, SVMs were used for toxicity prediction [20]. We imagine that our approach could also be used to detect the impact of particular genetic mutations, given earlier work suggesting that there may be significant differences in network activity [21].

## Conclusions

We report key differences in the developmental spontaneous activity patterns of cultured networks of hippocampal and cortical neurons. We have proposed a quantitative framework for evaluating these patterns. Our database of recordings and computer programs are all freely available for others to build upon. Future work in this area could be to dissect the cellular or network mechanisms driving the differences in network activity. For example, the differences between the cortical and hippocampus cultures could reflect molecular differences in cells or synapse or cellular differences in the populations of cells. Alternatively, differences in functional connectivity might partially account for these results [7]. Dissecting these differences will require a detailed understanding on the diversity of cell types defined by single cell transcriptomes in these brain regions, which is still lacking.

## Materials and Methods

### Primary neuronal culture

Primary cultures of dissociated hippocampal and cortical neurons were prepared from embryonic day (E) 17–18 mice. Hippocampi / cortices were dissected from E17.5 mouse embryos (2–4, pooled) and transferred to papain (10 units/mL Worthington, Lakewood, NJ) for 22 min at 37 °C. Cells were manually dispersed in Dulbecco’s Modified Eagle’s Medium containing 10% v/v foetal bovine serum and centrifuged twice at 400 g for 3.5 min. The final pellet was resuspended in Neurobasal/B27 supplemented with 0.5 mM Gln (Invitrogen), and dissociated cells (2 *×*10^5^ per dish) were seeded in the centre of poly-D-lysine / laminin coated multielectrode arrays (60MEA200/30-Ti, Multi Channel Systems, Reutlingen, Germany) containing 600 μl full Neurobasal medium. Zero-evaporation lids [22] were fitted and the MEAs housed in tissue culture incubators maintained humidified at 37 °C and 5% CO_2_ / 95% air. Twenty-four hours post-plating, sample MEAs were placed on an inverted microscope with heated stage (Axiovert 200; Zeiss) and photographed through a 32x phase objective at five different fields of view. These images were then analysed with CellProfiler [23] to quantify the neuronal density over the electrode array, giving an average value of 1500 cells/mm^2^.

At 3–4 days *in vitro*, cultures were fed by replacing 200 μl medium with pre-warmed fresh full Neurobasal medium. Cultures were subsequently fed using the same method after each recording, equating to a one-third medium change twice per week.

All procedures were performed in accordance with the United Kingdom Animals (Scientific Procedures) Act 1986. The mouse line used in this study was C57BL/6-*Tyr*^*c-Brd*^ (C57; albino C57BL/6).

### MEA recording

Multielectrode arrays and all data acquisition hardware and software were from MultiChannel Systems (Reutlingen, Germany). Pairs of MEAs were interfaced with duplex 60 channel amplifiers and 15 minute recordings of spontaneous action potentials were made twice per week during the four weeks following plating. MEAs were heated and kept under a light flow of 5% CO_2_ / 95% air during recordings. Signals were digitised with a 128-channel analogue/digital converter card at a rate of 25 kHz and filtered (100 Hz High pass) to remove low frequency events and baseline fluctuations. Action potentials were detected by crossing of threshold set to a fixed level of -20 μV, which typically approximated to 6–8 standard deviations from the baseline noise level. Record samples (1 ms pre- and 2 ms post-crossing of threshold) confirmed the characteristic action potential waveform. Application of tetrodotoxin (1 μM) totally abolished spiking activity, confirming the absence of false positive event detection using these methods. Spikes were not sorted to distinguish signals generated by individual neurons, so represent multiunit activity. Action potential timestamps were extracted using batch scripts written for NeuroExplorer [24] and analysed using software developed in the R statistical programming environment to compute parameters that quantitatively describe network activity. In total, 214 recordings were taken from 32 arrays of cultured cortical neurons, and 329 recordings from 61 arrays of cultured hippocampal neurons.

### Data analysis

To summarise a fifteen minute recording of network activity, we computed the following features. As all recordings detected activity from multiple electrodes, we calculated summary scalar values (termed the “array value” below) by summarising the information from multiple electrodes. In this way, each recording was then represented as an eleven-dimensional vector.

1. **Firing rate**. The mean firing rate of each electrode was calculated. The array value was the median of all electrode firing rates.
2. **Within-burst firing rate**. Bursts were detected independently on each electrode using our implementation of the Max Interval method from Neuroexplorer [24]. The parameters for burst detection were: maximum beginning ISI 0.1 s; maximum end ISI — 0.25 s; minimum interburst interval — 0.8 s; minimum burst duration — 0.05 s; minimum number of spikes in a burst — 6. For each electrode we calculated the mean of the firing rate during each burst. The array value was the median of the within-burst firing rates, ignoring electrodes where no bursts were detected.
3. **Burst rate**. For each electrode we calculated the number of bursts per minute. The array value was as per feature 2.
4. **Burst duration**. The electrode value was the mean duration of bursts on that electrode. The array value was as per feature 2.
5. **Fraction of spikes in bursts**. The electrode value was the total number of spikes classified as belonging to a burst divided by the total number of spikes on the electrode. The array value was as per feature 2.
6. **CV of IBI**. The electrode value was the coefficient of variation (s.d. divided by mean) of the interburst intervals. The array value was as per feature 2.
7. **Rate of network spikes**. Network spikes were defined as the array-wide average population activity [10]. It is defined as dividing time into small bins (here 3 ms) and counting the number of electrodes that generated at least one action potential in that bin. A network spike is then defined as the period of time when more than a threshold (here n=10) electrodes are simultaneously active. The array value was the number of network spikes per minute.
8. **Network spike peak**. During each network spike we found the maximum number of active electrodes. The array value was the median of the values from each network spike in a recording.
9. **Network spike duration**. The duration of each network spike was the time (in seconds) that the count of active electrodes exceeded the threshold value. The array value was as per feature 8.
10. **Mean correlation**. Given two different spike trains from the recording, we calculated the correlation between them using the spike time tiling coefficient [11] with the coincidence window of Δ*t* = 5 ms. (We also tried Δ*t* = 50 ms and 0.5 ms, but results were qualitatively similar.) The array value was the mean of all distinct pairs of electrodes.
11. **Fraction of electrodes exhibiting theta bursting**. For each electrode, the log interspike interval histogram was calculated and smoothed with the default kernel density estimation routine in R. A spike train was classed as showing theta bursting if a peak was present in the 4–10 Hz band of the histogram. The array value was the fraction of electrodes on the array that were classified as theta bursting.

The Mann-Whitney test was used to test whether the median array values at any given developmental age differed between the hippocampal and cortical networks. The p values were then corrected for multiple comparisons using the false discovery rate method [25].

### Clustering and classification

Standard principal components analysis was performed (with variance normalisation for each feature) for all feature vectors of any given age. Two standard machine learning classifiers were tested: classification trees with boosting (“random forests”) and support vector machines (SVMs) using radial kernel functions with γ = 1/11 [15]. For each age, we built binary classifiers to predict the region (CTX/HPC) based upon the eleven features measured from each recording. For both classifiers, we used 2/3 of the recordings as training data, with the remaining 1/3 used as test data. Performance is reported as mean percentage of correct classifications, averaged over 500 repeats using different splits of the data into training and test sets. The classification tree approach allows us to assess the relative importance of features by measuring the degree to which they decrease the Gini index [15, p319]. These values were normalised to the value of the top-performing feature.

### Data and code availability

Statistical analysis was performed in the R programming environment using the SJEMEA package [26]. Data files containing the spike times from all recordings analysed here were stored in the HDF5 format using the framework created for spontaneous activity in retinal recordings [27]. The only addition to the framework was a new metadata item `/meta/region` containing either the phrase “CTX” or “HPC” depending on the network type. All data files and analysis code relating to this paper are freely available at http://github.com/sje30/g2chvc. This includes all the material required to regenerate the figures and Table in this article.

## Abbreviations

CTX: cortex
DIV: days *in vitro*
HPC: hippocampus
MEA: multielectrode array
SVM: support vector machine

## Acknowledgements

### Competing interests

The authors declare that they have no competing interests.

### Authors’ contributions

PC, AM and SGNG conceived and designed the project. PC and AM performed all the experiments. EC and SJE provided analysis tools. PC, EC and SJE analysed the data. SJE drafted the manuscript; PC, EC, SGNG and SJE edited the manuscript. All authors read and approved the final version of the manuscript.

### Acknowledgements

Thanks to Diana Hall, Johannes Hjorth and Ole Paulsen for comments on this work. PC and AM were supported by the Wellcome Trust Genes to Cognition programme. PC received additional support from the BBSRC (BB/H008608/1). EC was supported by a Wellcome Trust PhD studentship and Cambridge Biomedical Research Centre studentship. SJE was supported by an and EPSRC grant (EP/E002331/1).

